# Neuroligin dependence of pharyngeal pumping reveals an extrapharyngeal modulation of *Caenorhabditis elegans* feeding

**DOI:** 10.1101/341982

**Authors:** Fernando Calahorro, Francesca Keefe, James Dillon, Lindy Holden-Dye, Vincent O’Connor

## Abstract

The integration of distinct sensory modalities is essential for behavioural decision making. In *C. elegans* this process is coordinated by neural circuits that integrate sensory cues from the environment to generate an appropriate behaviour at the appropriate output muscles. Food is a multimodal cue that impacts on the microcircuits to modulating feeding and foraging drivers at the level of the pharyngeal and body wall muscle respectively. When food triggers an upregulation in pharyngeal pumping it allows the effective ingestion of food. Here we show that a *C*. *elegans* mutant in the single orthologous gene of human neuroligins, *nlg-1* are defective in food induced pumping. This is not explained by an inability to sense food, as *nlg-1* mutants are not defective in chemotaxis towards bacteria. In addition, we show that neuroligin is widely expressed in the nervous system including AIY, ADE, ALA, URX and HSN neurones. Interestingly, despite the deficit in pharyngeal pumping neuroligin is not expressed within the pharyngeal neuromuscular network, which suggests an extrapharyngeal regulation of this circuit. We resolve electrophysiologically the neuroligin contribution to the pharyngeal circuit by mimicking a food-dependent pumping, and show that the *nlg-1* phenotype is similar to mutants impaired in GABAergic and/or glutamatergic signalling. We suggest that neuroligin organizes extrapharyngeal circuits that regulate the pharynx. These observations based on the molecular and cellular determinants of feeding are consistent with the emerging role of neuroligin in discretely impacting functional circuits underpinning complex behaviours.

## INTRODUCTION

Complex cognitive function is underpinned by discrete microcircuits that are integrated to control behaviour in a manner appropriate for the environmental context (LeDoux 2000, Barkus, McHugh et al. 2010, O’Connor, Finger et al. 2010). Arguably one of the most important environmental stimuli for an animal is its food. This relationship has been subjected to extensive investigation for the model genetic animal *Caenorhabitis elegans* with a view to providing insight into the fundamental behavioural relationship of an organism with its food source (Shtonda and Avery 2006, Greene, Brown et al. 2016, Thutupalli, Uppaluri et al. 2017). For *C. elegans* this food is bacteria which it ingests by increasing its rate of pharyngeal pumping, a process that is controlled by pharyngeal motor neurons that direct coordinated cycles of pharyngeal radial muscle contraction and relaxation (Franks, Holden-Dye et al. 2006). Intrinsic control of the pharynx modulated by MC and M3 neurons is required for an efficient intake of bacteria (Avery 1993, Avery 1993a). The pharyngeal circuit connects to the nervous system via a single neuron RIP which provides a route for extrapharyngeal control, however in addition the pharyngeal system is regulated by neurohumoral signalling, as ablating RIP neuron results in essentially normal pharyngeal activity to changing food context (Albertson and Thomson 1976, Dalliere, Bhatla et al. 2016). These mechanisms provide a means whereby the rate of pharyngeal pumping is influenced by the presence of food (Horvitz, Chalfie et al. 1982, Rogers, Franks et al. 2001).

MC neurons act as pacemakers to control the rapid activation of the pumping rate in *C. elegans* (Avery and Horvitz 1989, Raizen and Avery 1994). This is facilitated by the motorneuron M3 causes repolarization of the muscles via coordinated glutamate release. This relaxation of the metacorpus and isthmus is critical in trapping the bacteria in the corpus (Albertson and Thomson 1976). Electrophysiological recording of the pharyngeal neural network via electropharyngeogram (EPG), provides a measurement of fine features that underpin pumping (Avery, Raizen et al. 1995). This allows one to record the electrophysiological activity of both M3 and MC neurons, as well as a characterization of synaptically driven events from both excitatory and inhibitory neurones. This has identified neuronal signalling and neuromodulatory components of the pharyngeal microcircuit (Raizen and Avery 1994, Dillon, Andrianakis et al. 2009). Two serotonergic neurons play a key role in mediating the pharyngeal response to the availability of food, the pharyngeal secretory neuron NSM and the chemosensory neuron ASD (Li, Li et al. 2012). The neuron NSM is embedded within the pharyngeal circuit and implicated in the acute response to newly encountered food (Iwanir, Brown et al. 2016). Although functional NSM neurons are required for bursts of fast pumping in the presence of food, serotonin production in HSN neurons is sufficient to trigger food-dependent pumping dynamics when serotonin production in NSM is perturbed. This suggests that the HSN neurons, located in the middle of the worm body and send no processes into the pharynx, are involved in the regulation of feeding (Lee, Iwanir et al. 2017). Finally, the role played by ADF in feeding regulation is related to sensing fluctuations in the environment (Lee, Iwanir et al. 2017).

In addition food availability also modifies the worm’s locomotory behaviour and induces high probability of dwelling. This behaviour promotes the worm residence on a patch of food. This dwelling behaviour is interspersed with roaming, a behaviour that allows the worms to explore their environment (Flavell, Pokala et al. 2013). Thus, *C. elegans* exhibits a repertoire of behavioural responses to food that requires coordination of precisely organised neural circuits. How the upstream sensory inputs are integrated and modulated to bring about this coordinated behavioural response is important in understanding the systems level control of foraging and feeding.

In this study we have focused on the role of neuroligin in the regulation of this food-dependent behaviour. Neuroligin is a synaptic protein intimately involved in molecular organization of discrete circuits (Calahorro 2014, Bemben, Shipman et al. 2015). In particular, in its absence there is an impairment in the organization of the excitatory-inhibitory balance in neuronal circuits (Varoqueaux, Aramuni et al. 2006). *nlg-1* is the *C*. *elegans* orthologue of human *NLGN1* presents at most of synapses, and with a conserved organization in functional domains (Calahorro 2014). *C. elegans nlg-1* mutants have previously been shown to express a range of sensory deficits (Calahorro, Alejandre et al. 2009, Hunter, Mullen et al. 2010, Calahorro and Ruiz-Rubio 2012). Indeed, *C. elegans* provide an excellent opportunity to delineate the role of neuroligin in the functional organisation of sensory driven behaviours as they express one orthologous of neuroligin, *nlg-1.* In *C*. *elegans nlg-1* is selectively expressed in a subset of neurons. Here, we provide evidence that neuroligin functions in an extrapharyngeal circuit that coordinates a feeding response by modulating the cue promoting the pharyngeal function. This further supports its fundamental role in the integration of sensory information and co-ordination of activity via discrete microcircuits.

## MATERIAL AND METHODS

### Culturing of *C. elegans* and strains used

*C. elegans* strains were maintained under standard conditions (Brenner 1974, Molin, Schnabel et al. 1999). Worms were synchronized by picking L4 larval stage to new plates 18h prior to performing behavioural experiments. The strains used were: *nlg-1* (*ok259*) *X* (6x outcrossed); *nlg-1* (*ok259*) X, Ex [pPD95.77 (P*nlg-1::nlg-1* Δ#14); P*myo-3::gfp*]; *nlg-1* (*ok259*) X, Ex [pPD95.77; P*myo-3::gfp*]; RM371, *pha-1(e2123*), sIs [P*nlg-1*::*yfp* + pCI(*pha-1*)]; MT6308 *eat-4* (*ky5*) *III* (2x outcrossed); CB156, *unc-25 (e156) III* (1x outcrossed); CB382, *unc-49 (e382) III* (2x outcrossed).

We identified neurons that express *nlg-1* reporter constructs based on their cell position and, co-labelling neurons with mCherry/RFP/DsRed-based reporters.

The co-labelled reporter transgenes were:

- oyls 51, aka: (P*srh-142*::*rfp*). A specific marker for ADF chemosensory neuron (Xu, Choi et al. 2015).
- oyIs44, oyIs44 [P*odr-1*::*rfp* + *lin-15*(+)]. A specific marker for AWB, AWB odorsensory neurons (Sarafi-Reinach, Melkman et al. 2001)
- ofEx205 [pBHL98 (*lin-15*ab+); P*trx-1*::*DsRed*]. A specific marker for ASJ sensory neuron (Miranda-Vizuete, Fierro Gonzalez et al. 2006).
- dbEx719 [P*npr-5*::mCherry + P*unc-122*::*gfp*]. A specific marker for sensory neurons: ADF, ASE, ASG, ASI, ASJ, ASK, AWA, AWB, IL2; interneurons: AIA, AUA; phasmids sensory neurons: PHA, PHB (Cohen, Reale et al. 2009).
- sIs [P*eat-4*::m*rfp*]. A specific marker for neurons expressing the vesicular glutamate transporter, *eat-4* (Serrano-Saiz, Poole et al. 2013).
- otIs151 [P*ceh-36*::*rfp* + *rol-6* (su1006)]. A specific marker for chemosensory neurons: AWC and ASE (Chang, Johnston et al. 2003).
- otIs181 [P*dat-1*::*mCherry* + P*ttx-3*::*mCherry*]. Specific markers for both dopaminergic neurons (P*dat-1*): ADE, PDE, CEP; and for the sensory interneuron AIY (P*ttx-3*) (Flames and Hobert 2009).
- vsIs108 [P*thp-1*::*rfp* + P*unc-13*::*gfp* + *lin-15*(+)]. A specific marker for HSM and NSM serotonergic neurons (Tanis, Moresco et al. 2008).

### Cloning and transgenic methods

#### cDNA cloning of the C. elegans nlg-1 gene

The genomic sequence of *nlg-1* was used to design the following specific primers: *nlg-1*forward 5′-GGCAT***ggatcc***CATTTATCTTCTTCTCC-3′ and *nlg-1*reverse 5′-GTA***gaattc***GTTAGACCTGTATCTCTTCC-3′, to amplify by PCR the coding region corresponding to the Δ#14 transcript, the dominant neuroligin isoform in the adult stage (Calahorro, Holden-Dye et al. 2015), from the cDNA clone yk1657a10 (from Yuji Kohara, National Institute of Genetics, Mishima, Japan).

#### Rescue constructs and transgenic methods

The 2.5 kb promoter region of *nlg-1* was amplified using the P*nlg-1*forward 5′-T***tctaga***CATATTTTTGGGGAGGCTTTC-3′ and P*nlg-1* reverse 5′-GAAGGAGAAGAAGATAAATG***ggatcc***ATGC-3′ primers from the C40C9 cosmid clone (from Sanger Institute, UK). The promoter sequence was cloned using the *XbaI*/*BamHI* site in a pDD95.77 vector where the GFP was removed. Finally, the *nlg-1* Δ#14 sequence was fused to the *nlg-1* promoter using the *BamHI/EcoRI* sites. The promoter was authenticated by sequencing using the following primers: 5′-AAGCTTGCATGCCTGCAGGTCGAC-3′, 5′-AAGCTTGCATGCCTGCAGGTCGAC-3′; and the following primers for *nlg-1*: 5′-AATGCAGACTGGAGAAACTTTG-3′, 5′-CTATTACCAGAGCAAGACGATG3′, 5′-GCTTCTCTGGTTTCTCTTCTTATG-3′, 5′-CTGTTTCCTTTCCATTCTTGTGC-3′, 5′- AGAATGGAAAGGAAACAGAGCC-3′, 5′-GTGCGATGCGGATAGTAAGGG-3′.

#### Transgenic methods

*nlg-1* (o*k259*) *X* one day old adults were microinjected with *nlg-1* plasmid (prepared as described above; 50 ng/μl) together with the “marker” plasmid P*myo-3::gfp* (30 ng/μl) as previously described (Mello & Fire 1995). For controls, *nlg-1* (o*k259*) *X* animals were microinjected with “empty rescue” plasmid and the “marker plasmid” P*myo-3::gfp* at the same concentration used for generating the rescue transgenic lines. Microinjection mixes were prepared using double-distilled water.

### Behavioural assays

#### Feeding behaviour

Feeding behaviour was visually scored by counting the number of pharyngeal pumps for 1 min using a binocular dissecting microscope (x63). A single pharyngeal pump was defined as one contraction-relaxation cycle of the terminal bulb of the pharyngeal muscle.

To count pharyngeal pumps on food, one day old adult worms (L4+1) grown at 20 °C, in standard NGM-*E. Coli* OP50 plates, were gently picked onto the middle of the bacterial lawn (OD_600_ of 0.8 AU seeded the day before). After 10 mins the pharyngeal pumping was recorded using a hand counter. Five consecutive measurements (1 min each) were made and the mean of pharyngeal pumping rate for this time period was then calculated.

To count pharyngeal pumps off food, one day old adult worms grown at 20 °C, in standard NGM-*E. Coli* OP50 plates, were gently picked onto the middle of a 9 cm diameter non-food plate and left for 5 min to clean itself of attached bacteria. Animals were finally transferred onto a second clean non-food plate, where pumping rate was scored after 10 mins. Five consecutive measurements (1 min each) were made and the mean of pharyngeal pumping rate for this time period was then calculated. During this experiment, worms dried out after migration off the edge of the agar plate were censured.

#### Chemotaxis; Food race assays

9 cm NGM plates were poured the day before the assay and kept overnight at room temperature. These plates were seeded with a spot of 50 µl *E. coli* OP50 at an optical density of 0.8 AU (OD_600nm_) displaced 2 cm from the edge of the plate and incubated overnight. The day before the assay 50-100 L4 animals were picked onto NGM *E*.*coli* OP50 plates and kept at 20˚C for 18 h. These L4+1 worms were washed twice in M9 buffer to remove residual bacteria adhered to the body. The washed worms were added in a minimal volume 2 cm from the edge opposite to the food patch. The number of animals reaching food was recorded every 10 min for of time of 2 h. The cumulative number that reached the food spot was calculated.

### Electropharyngeogram (EPG) Recordings

#### Recording the activity on the pharyngeal neural network

The extracellular electrophysiological activity in the pharyngeal network was measured using the microfluidic device Neurochip as previously described (Hu, Dillon et al. 2013). L4+1 worms were loaded into the NeuroChip in Dent’s saline (in mM: 10 d-glucose, 140 NaCl, 1 MgCl_2_, 3 CaCl_2_, 6 KCl, and 10 HEPES; pH 7.4) supplemented with 0.01% BSA (w/v). Extracellular voltage recordings were made in “bridge” mode, and the extracellular potential was set to 0 mV using the voltage offset immediately prior to recording. Data were acquired using Axoscope (Axon Instruments) and stored for subsequent offline analysis. EPG traces were annotated with AutoEPG software (Dillon, Andrianakis et al. 2009). Statistical analysis was performed using one way ANOVA with Bonferoni multiple comparisons post-test. Recordings were made in both Dent’s alone or when loaded into the device in Dent’s containing 5 mM 5HT (5-Hydroxytryptamine **-** serotonin creatinine sulfate monohydrate used for electrophysiology experiments was obtained from Sigma-Aldrich ^®^). EPGs were recorded for 3-5 min from when the animal was trapped in the recording microchannel, conditions under which the 5 mM serotonin induced pumping (Hu, Dillon et al. 2013). The electropharyngeograms (EPGs) recorded single pump corresponds to the coordinated contraction and relaxation of the pharynx. Recordings were analysed offline using a signal peak detection and measurements made to define: frequency; average duration of single EPGs; number of *P* waves per EPG, regulated by activity of the inhibitory motorneurone M3; shape of the EPG using the *R* to *E* ratio, closely associated with muscle depolarization (contraction) at the beginning of a pump and repolarization (relaxation) at the end of a pump; pattern of activity, under neural control.

### Imaging

#### Imaging worms

A Nikon Eclipse (E800) microscope was used to image fluorescence from cells expressing GFP or RFP proteins under cell specific promoters. Worms were put into 0.5 μL M9 buffer containing 25 mM sodium azide on a thin 2% agarose pad. Immobilized worms were covered with a 24×24 mm cover slip, and viewed with 40–63X objective magnification for no more than 10 minutes after addition of levamisole. At least 10 independent worms per strain were analysed. The position of cell bodies relative to neuronal ganglia, as well as the dominant visible neuronal process were imaged by both DIC optical and epifluorescence. Images were acquired through a Hamamatsu Photonics camera software, and were cropped to size, assembled, and processed using Abode PhotoShop^®^ (Adobe Systems) and ImageJ (NIH) softwares.

#### DiI staining

A stock solution of 2mg/ml DiI in dimethyl formamide was prepared, and then a 1:200 dilution in M9 was used for staining. Animals were transferred into a microtiter well containing 150 µl stain and incubated 2-3 hours in dark and room temperature. Using a pipette the animals were transferred to a fresh plate and left crawl on the bacterial lawn for about 1 hour to destain. Finally, animals were immobilized on an agar pad with sodium azide and visualized by fluorescence.

#### Pharynx dissection

Around 10 L4+1 animals were transferred into a 3cm Petri dish containing 2mL of M9 supplemented with BSA and incubated for 1 h at 4°C to immobilize the animals. Then animals were observed under a normal dissection microscope (40x magnifications). Dissection was performed as previously described (Franks, Murray et al. 2009). The isolated pharynxes were pipetted onto a 2% agarose pad and covered with a cover slip prior to experimental observation. A total of five isolated pharynxes were checked.

### Statistical analysis

Data are expressed as mean ±SEM and were analysed using Graphpad Prism version 7.00 (GraphPad Software Inc.) using unpaired Student’s t-test, one-way ANOVA (Analysis of Variance), or two-way ANOVA as indicated in figure legends. For ANOVA the data were subjected to post-hoc analysis using the method indicated in the figure legends. Significance level was set at p<0.05.

## RESULTS

### Neuroligin is required to upregulate feeding in the presence of bacteria

In a comparison of pharyngeal pumping of N2 (wt) (244±7 pump.min^−1^) the *nlg*-1 mutants showed a reduction of ≈26% (180±12pump.min^−1^) (**Fig 1A**). This was selective to the on food context, as the pharyngeal pumping rate of *nlg-1* mutants and wild-type animals was not significantly different 15 mins after transferring to a non-food plate (39±4 min^−1^ for wild-type and 38±4 min^−1^ for *nlg-1*; n=18 animals for each strain; p>0.05. Data not shown). This food context dependent reduction in pharyngeal pumping in *nlg-1* was rescued by expressing a cDNA encoding the *nlg-1* Δ#14 isoform from the *nlg-1* promoter. The transcript Δ#14 is the dominant isoform in the adult stage (Calahorro, Holden-Dye et al. 2015). This fully rescued the pharyngeal deficit in *nlg-1* mutants **(Fig 1A).** These results indicate a selective role for NLG-1 dependent signalling in maintaining a high rate of pharyngeal pumping in the presence of food. It is unlikely that this deficit can be explained by an inability of *nlg-1* to sense food they show chemotaxis towards a point source of bacteria at the same rate as wild-type worms (**Fig 1B**).

**Figure 1.**
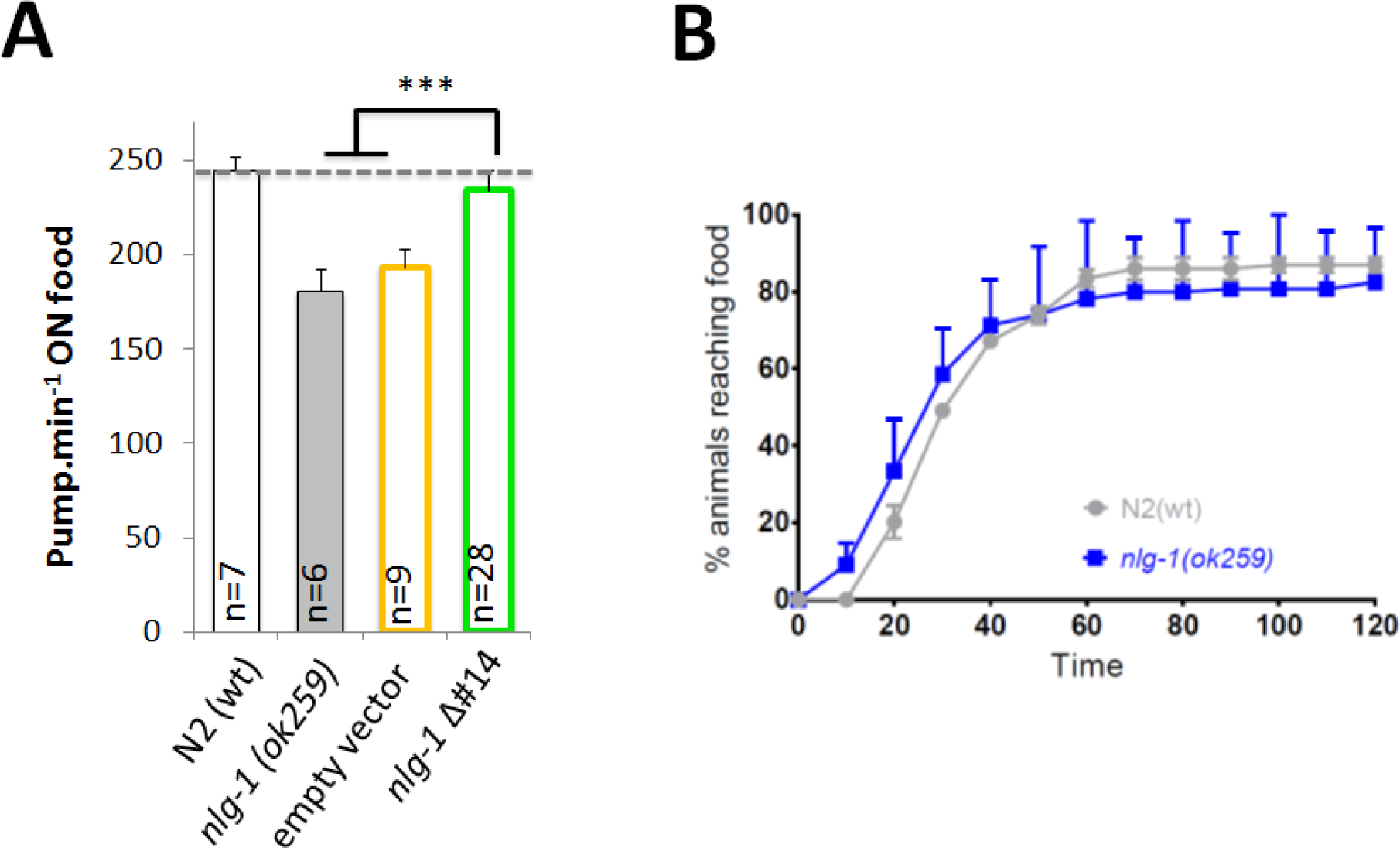
**A. Neuroligin deficient mutants have lower pharyngeal pumping rates on food** Nematodes at the L4 stage were placed onto fresh seeded (with *E. coli* OP50) plates, and left overnight before the assay was performed. Each nematode was observed for 5 mins and the number of pharyngeal pumps within each min was recorded. The bars represent the mean pharyngeal pumping rate ±SEM. The means were calculated from data collected from repeats of the same experiment conducted on different days, giving the specified ‘n’ number, but in each case the experiment was paired with both N2(wt) and *nlg-1 (ok259)*. Rescue plasmid *Pnlg-1*::*nlg-1 Δ#14* was co-injected with the marker plasmid *Pmyo-3*::*gfp.* To generate control lines the marker plasmid *Pmyo-3::gfp* was co-injected together with the empty plasmid used to build the rescue plasmids. Data are expressed as the mean ± SEM: 244 ±7 min^−1^ for N2 (n=7), 180 ±12 min^−1^ for *nlg-1 (ok259)* (n=6), 192±10 min^−1^ for control line (n=9) and 233 ±10 min^−1^ for human *NLGN-1* rescue line (n=28). The data were subjected to one-way ANOVA followed by a Bonferroni’s post hoc test. There was a significant difference between *nlg-1 (ok259)* and the control line means compared to the human *nlg-1* rescue line (***P ≤0.001). No significant difference was found between the means of *nlg-1 (ok259)* and the control line (ns, P>0.05). **B. *nlg-1* mutants are not chemotaxis deficient for food** Each “food race” was performed for a population of animals (50-100 per race) on 9 cm agar plates with a single spot of *E. coli* OP50 2 cm from the edge of the plate. At time zero, staged worms (L4 plus 1 day) were aliquoted on to the plate diametrically opposite the food source. Every 10 min, the number of worms that had reached the food was counted and expressed as a percentage of the total population. Data are expressed as the mean ±SEM of ‘n’ experiments where each experiment is a single food race. Statistical analysis was performed using Student’s unpaired t-test. There was no significant differences between N2(wt) and *nlg-1* mutants at every single time point (n=3). t=10, p=0.24; t=20, p=0.603; t=30, p=0.706; t=40, p=0.857; t=50, p=0.999; t=60, p=0.819; t=70, p=0.776; t=80, p=0.776; t=90, p=0.813; t=100, p=0.779; t=110, p=0.779; t=120, p=0.824.

### NLG-1, GABA and glutamate signalling act as positive modulators of feeding

Experiments in which the RIP neuron is ablated indicate there may be important extrapharyngeal determinants that drive the pharyngeal microcircuit via volume transmission. However, these RIP ablations may also highlight a clear potential for intrinsic signalling pathways embedded within the pharyngeal nervous system to act as regulators of the food induced pumping (Dalliere, Bhatla et al. 2016). Insight into the contribution of distinct signalling pathways to pharyngeal regulation can be obtained from analysis of an electrophysiological recording called the electropharyngeogram, EPG (Avery, Raizen et al. 1995, Cook, Franks et al. 2006, Franks, Holden-Dye et al. 2006) and by comparing the EPG waveform between wild-type and mutants that are defective in the signalling pathways of interest. Therefore, we made EPG recordings in order to provide insight into the neural mechanisms underpinning the *nlg-1* dependent pharyngeal behaviour. Each EPG waveform represents the activity of the pharyngeal neuromuscular circuit during a single contraction-relaxation cycle of the pharyngeal muscle that underlies a pharyngeal pump and carries information about the activity of specific neurons that regulate pump frequency, pump duration and the discrete functional organisation of each pharyngeal pump (Franks, Holden-Dye et al. 2006). The typical EPG harbours potentials that relate to the MC synaptic activity (‘e’), the synchronous contraction of the muscle (‘E’), the inhibitory signals from M3 (‘P’), and the rapid relaxation of the muscle (‘R’) and the repolarization of the terminal bulb (‘r’) (**Fig 2Ai**).

**Figure 2.**
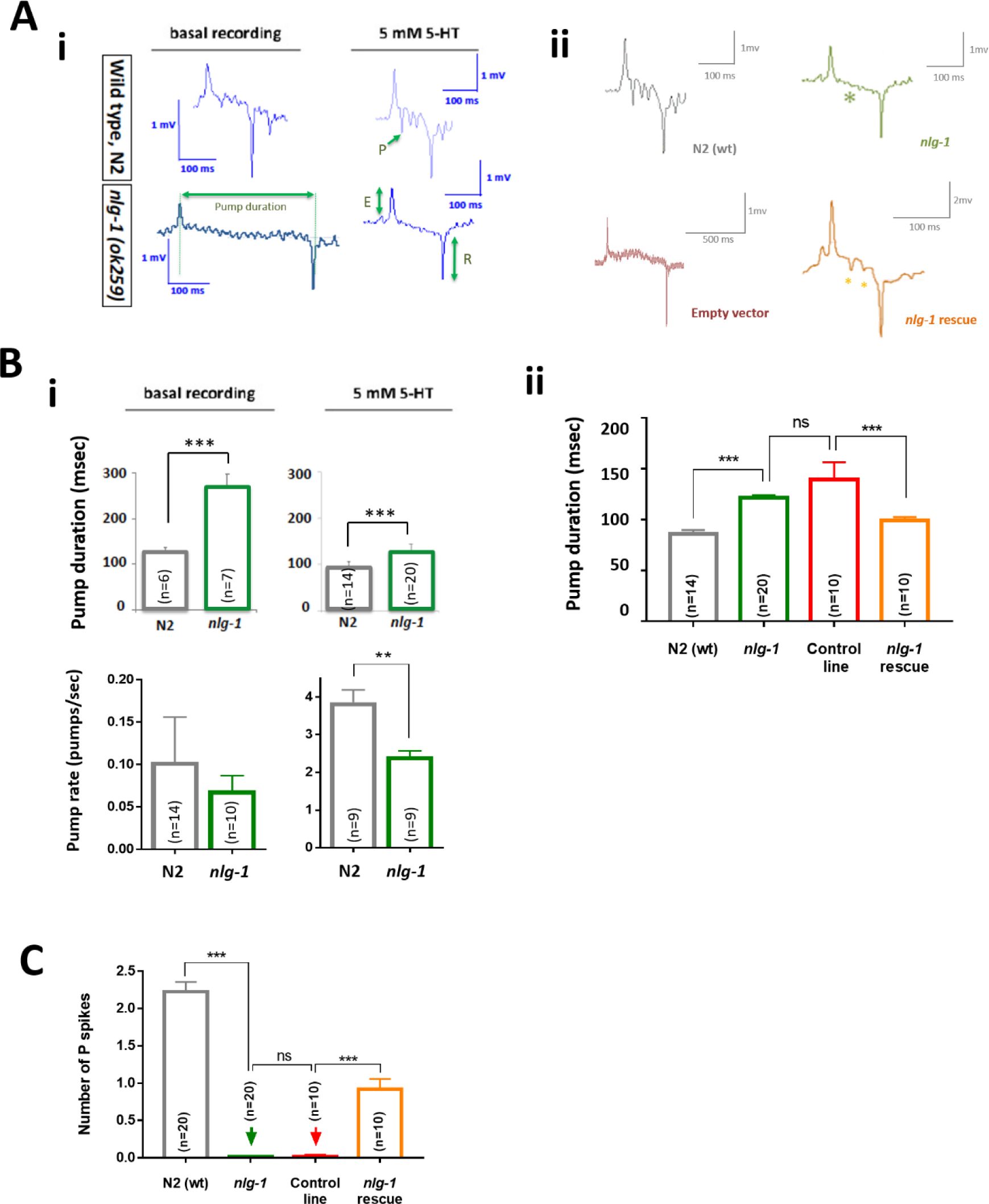
NLG-1 modulates the contraction-relaxation cycle of the pharyngeal neuromuscular system that underpins feeding. **A(i)**. Representative unfiltered electropharyngeogram (EPG) traces recorded using the NeuroChip with whole *C. elegans*, comparing the EPG waveform from both the wild-type and *nlg-1* in the presence or absence of 5-HT. Excitation [E], relaxation [R], and P spikes [P] are annotated in green, as well the pumping duration of an EPG; **A(ii)**. Representative unfiltered EPG signatures from N2 (wt) (grey), *nlg-1 (ok259)* (green), control empty transgene (red) and *nlg-1* rescue (yellow). The asterisks mark the trace to denote the absence or presence of *P* spikes in both *nlg-*1 and rescue transgenic strains respectively. **B(i)**: Neuroligin mutants have a longer pump duration and a subsequent decreased pump rate. Average of pump duration comparing N2 (wild-type) and *nlg-1 (ok259)* animals, in basal conditions (N2 (wt) 131 ±17 msec, *nlg-1* 271 ±28 msec), and pump rate (N2 (wt) 0.1 ±0.2 pump.sec^−1^, *nlg-1* 0.07 ±0.05 pump.sec^−1^). In the presence of 5mM 5-HT despite a comparable increase in pump rate (N2 (wt) 3.9 ±0.9 pump.sec-^1^, *nlg-1* 2.4 ±0.4 pump.sec^− 1^), the pump duration of the *nlg-1* mutant is longer (N2 (wt) 95 ±9msec, *nlg-1* 122 ±8 msec). **B(ii)**: Quantification of pump duration in N2(wt) and *ngl-1* mutant, comparing to control line (carrying the empty transgene) and *nlg-1* rescue line. Pump duration is reduced relative to control by introducing the *nlg-1 Δ#14* transgene version (N2 (wt) 86 ±4 msec, *nlg-1* 122 ±3 msec (***P ≤0.001). Control line 140 ±18 msec, *nlg-1* rescue 99 ±4 msec (***P ≤0.001)). **C.** *nlg-1* mutants show a significative reduction in the number of *P*s per EPG (2 ±0.1 N2 (wt) and 0 ±0 *nlg-1*; ***P ≤0.001), and it is partially restores in presence of the *nlg-1 Δ#14* transgene version (0 ±0 control line and 0.9 ±0.1 *nlg-1* rescue; ***P ≤0.001).

In the absence of 5-hydroxytryptamine (5-HT or serotonin), the frequency of EPGs is low for both wild-type and *nlg-1* (N2 (wt) 6.19±1.65 pump.min^−1^, *nlg-1* 4.18±1.32 pump.min^−1^; p>0.05) reflecting the fact that this is an ‘off-food’ pumping rate and therefore feeding rate is down-regulated (Dalliere, Bhatla et al. 2016). This is consistent with the low pump rate seen in animals off food. However, despite the similarity in EPG frequency between N2 and *nlg-1*, the EPG analysis revealed discrete changes in the EPG waveform when N2 (wt) and *nlg-1* mutants were compared: The *nlg-1* mutant exhibited a marked increase in pump duration (**Fig 2A-B**) and an absence of transient potentials called *P* waves that are dependent on the activity of the inhibitory glutamatergic neuron M3 (Avery 1993a) (**Fig 2C**). We next analysed the EPG waveform in the presence of 5-HT. This is a well-characterised positive regulator of pharyngeal pumping which is engaged in the presence of food (Niacaris and Avery 2003). In the presence of 5-HT EPG rate was elevated in both N2 (wt) and *nlg-1* (**Fig 2B**). Despite this 5-HT mediated increase in pumping, the *nlg-1* EPG waveform showed a significant reduction in the number of *P* waves, as well as an increased pump duration relative to recordings made from 5-HT treated N2 (wt) pharynxes (**Fig 2A**). The loss of these key signatures was rescued in the strain expressing *nlg-1* Δ#14 from the *nlg-1* promoter (**Fig 2B, C**).

To gain further insight into the neural basis of the effect of *nlg-1* on the EPG waveform we made a systematic comparison of EPGs in *nlg-1* compared to mutants of known regulators of EPGs. It is well established that the absence of *P* waves provides a readout of activity of the inhibitory glutamatergic motorneuron M3 (Avery 1993a) which regulates the duration of the pharyngeal contraction-relaxation cycle by accelerating muscle repolarisation. The loss of glutamate release from M3, as observed in the mutant *eat-4* (Raizen and Avery 1994), results in decreased *P* waves and increased pump duration (Raizen and Avery 1994). *nlg-1* mutants show a similar EPG signature to *eat-4* mutants (**Fig. 3**). This is consistent with an involvement of glutamate signalling that is intrinsic to the pharyngeal circuit as the increased pump duration and loss of *P* waves is characteristic of a role for the glutamatergic pharyngeal neuron M3.

**Figure 3.**
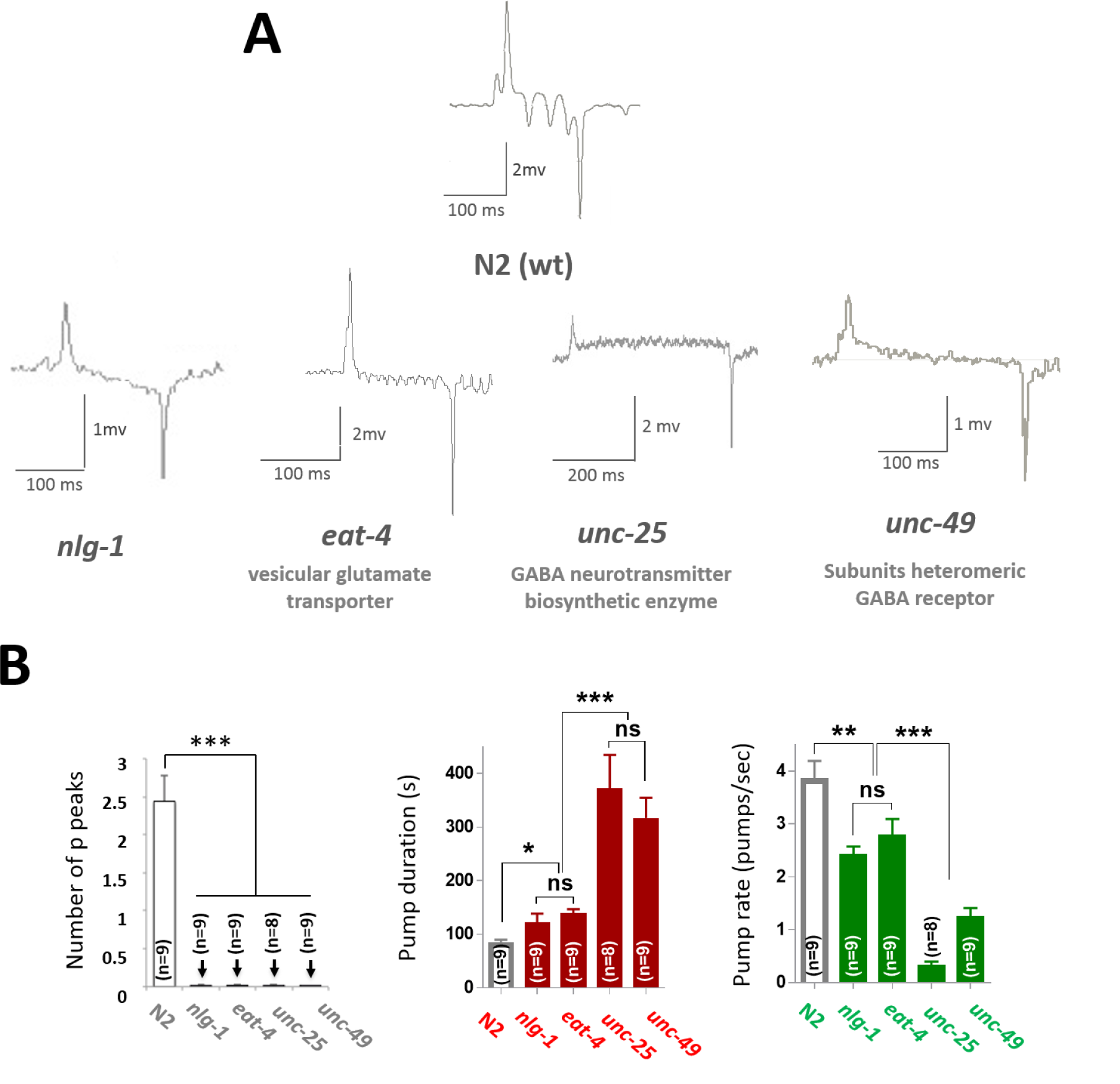
Glutamate and GABA signalling mutants exhibit a similar pharyngeal phenotype to *nlg-1*. **A.** Representative unfiltered electropharyngeogram (EPG) traces recorded using the NeuroChip comparing N2 (wt) signature with *nlg-1*, *eat-4*, *unc-25* and *unc-49* mutants. EPGs parameters comparison between neuroligin and GABA/glutamate mutants: *eat-4* (encodes a vesicular glutamate transporter), *unc-25* (encodes a GABA neurotransmitter biosynthetic enzyme), *unc-49* (encodes a subunit of a heteromeric GABA receptor). **B.** Realtive to N2 (wt), *nlg-1* (*P ≤0.05), *eat-4* (*P ≤0.05), *unc-25* (***P ≤0.001) and *unc-49* (***P ≤0.001) mutants show longer pump duration. In addition *nlg-1* (***P ≤0.001), *eat-4* (***P ≤0.001), *unc-25* (***P ≤0.001) and *unc-49* (***P ≤0.001) have reduced *P* spikes. Finally, both *unc-25* and *unc-49* mutans show a reduced pump rate relative to N2 (wt), *nlg-1* and *eat-*4 subsequent with the increase in pump duration. In basal conditions no EPGs were detected during 30 mins recordings for *unc-25* and/or *unc-49* mutants. These parameters were extracted from 2 min recording in presence of 5-HT (5mM). Statistical analysis was performed using one way ANOVA with Bonferroni multiple comparisons post-test.

Although GABA is not known to be a neurotransmitter within the pharyngeal microcircuit (Franks, Holden-Dye et al. 2006), GABAergic signalling has been reported to modulate pumping (Dalliere, Bhatla et al. 2016). Recent observations show a *nlg-1* dependence of GABAergic signalling in both *C. elegans* (Maro, Gao et al. 2015, Tu, Pinan-Lucarre et al. 2015) and mammals (Budreck and Scheiffele 2007, Fu and Vicini 2009). Motivated by these findings we next compared the EPG waveform for the GABAergic signalling mutants *unc-25* (*e156)* and *unc-49* (*e382*) with *nlg-1*. Surprisingly, given that neither GABA nor GABA receptors are present in the pharyngeal neurons, the EPG phenotype is even more marked in *unc-25* and *unc-49* (**Fig 3A, B**) than in *eat-4*, with a very extended pump duration and absence of *P* peaks. This therefore identifies a previously unreported GABAergic extrapharyngeal regulation of pharyngeal physiology. Moreover, the similarity in the EPG phenotype between the GABAergic mutants and *nlg-1* suggests neuroligin signalling might exert its organization of pharyngeal function at an extrapharygneal locus rather than at a site intrinsic to the pharyngeal circuitry. Therefore we next investigated the expression pattern of *nlg-1* in the pharyngeal and extrapharyngeal circuits.

### *nlg-1* is expressed in a subset of sensory neurons and interneurons

*nlg-1* is widely expressed in the nervous system of *C. elegans* including the head region (Hunter, Mullen et al. 2010). Although detailed expression has been ascribed to AIY, DAs, VAs, PVD, URA and an undefined subset of cholinergic and GABAeregic neurons (Hunter, Mullen et al. 2010, Hu, Hom et al. 2012, Maro, Gao et al. 2015, Tu, Pinan-Lucarre et al. 2015), these is no definitive information on expression in the pharyngeal microcircuit. Therefore, an important first step was to resolve *nlg-1* expression in the pharyngeal nervous system. This consists of 20 neurons that are embedded within the pharyngeal muscle and separated from the central nervous system of the worm by a basal lamina (Donna G. Albertson 1976). We dissected and isolated pharynxes from two *nlg-1* transcriptional reporter strains and established that there is no *nlg-1* expression in the pharynx or in its associated microcircuit (**Fig 4A**). Therefore neuroligin exerts its regulation of pharyngeal behaviour through extrapharyngeal circuits.

**Figure 4.**
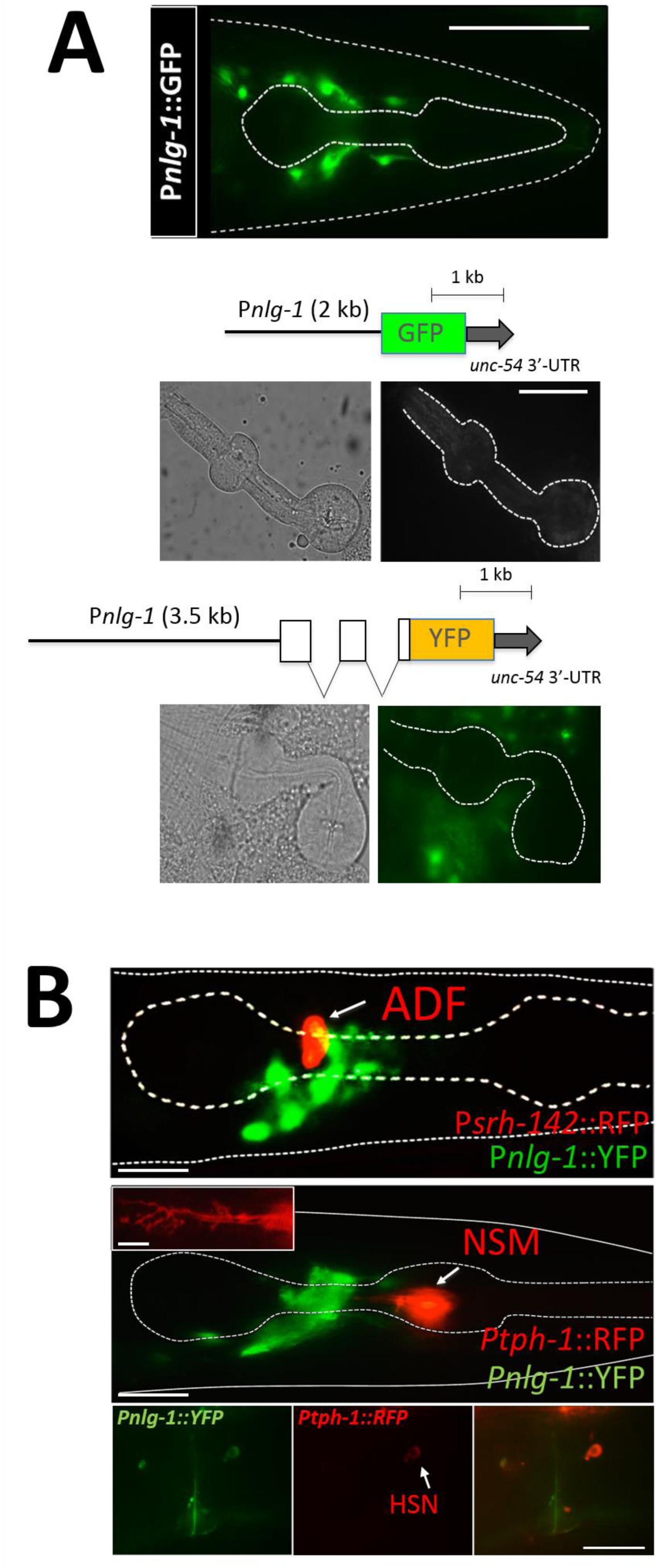
Neuroligin is expressed in a subset of extrapharyngeal neurons in the head ganglia. **A.** Neuroligin is expressed in a subset of neurons around the pharynx. An isolated pharynx from a transcriptional reporter strain shows non-expression of neuroligin in neurons of the pharynx. A cartoon of the characteristic endings of IL2 and URA cilia are shown. Two different transcriptional reporter strains were used to compare neuroligin expression level in pharyngeal neurons. The strain RM371 where the expression is driven by an integrated array carrying ^~^3.5 kb of *nlg-1* upstream sequence as well as regulatory elements present in the first exons of the neuroligin gene fused with YFP (upper panel), and the strain BC13535 where the expression is driven by an integrated array carrying ^~^2.9 kb upstream to the ATG fused with GFP (bottom panel). The cartoon above the isolated pharynx images shows detail of the transcriptional reporters for each strain. The scale bar indicates 50 μm. **B.** Neuroligin is not expressed in ADF and/or NSM neurons, the two major pharynx neurons modulating the feeding efficacy extrinsic and intrinsically respectively. However neuroligin is expressed in HSN neuron which triggers food-dependent pumping regulating the feeding extrapharyngeally. Images from co-localization with P*thp-1::rfp* show HSN expression around the vulva. A characteristic process to identify the NSM neuron is detailed in the box inset on the image. The scale bar indicates 50 μm.

To place *nlg-1* in the context of neural circuits that link food-related sensory inputs to pharyngeal behaviour (**Fig 1A**) we conducted detailed mapping of its expression pattern. Approximately 20 neurons express neuroligin in the anterior ganglia. We focused our attention on extrapharyngeal neurons that are known to regulate food-related behaviours (**Table 1**). To facilitate this, we carried out co-localization experiments using arrays expressing red fluorescent protein, RFP, in specific sensory neurons (**Table 1**). ADF is a sensory neuron which provides a serotonergic drive to increase pumping rate on food (Song, Faumont et al. 2013). We found no co-expression of neuroligin in ADF (**Fig 4B**) and no expression in major sensory neuron classes, using specific markers for the following neurons: ASJ, AWB, AWC and ASE neurons (**Fig 4, Supplement 1**). However, although neuroligin is not expressed in NSM we found expression in HSN neurons in the vulva area (**Fig 4B**). Finally, to further investigate expression in a subset of sensory neurons we tried to identify neuroligin expression in the DiI labelled amphid neurons: ADL, ASH, ASI, ASJ, ASK, AWB: Again we found no neuroligin expression (**Fig 5A**).

**Figure 5.**
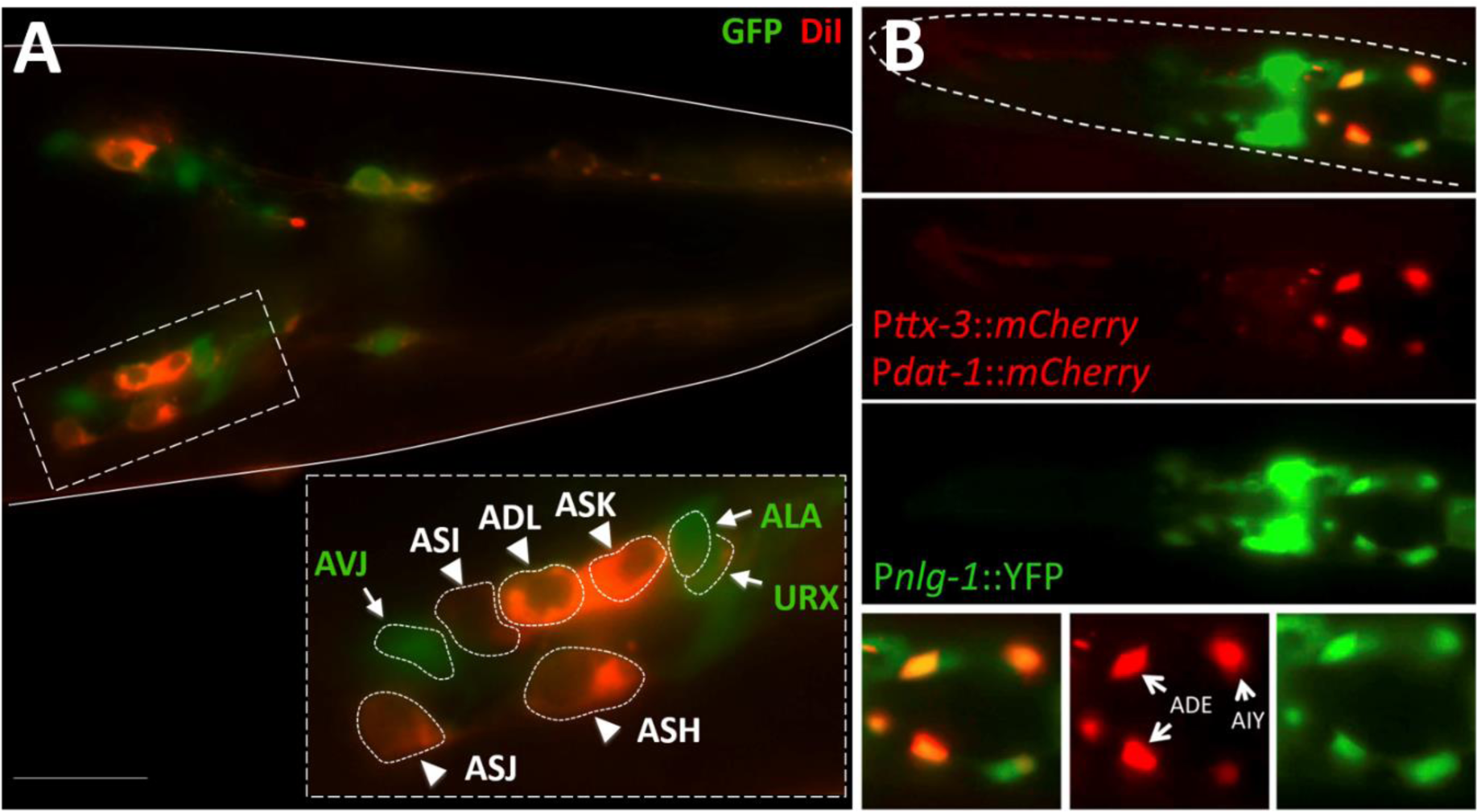
Neuroligin is expressed in a subset of sensory neurons and interneurons. **A**. A representative image from DiI staining of the strain carrying the P*nlg-1*::*yfp* array. The green signal corresponds to cells expressing YFP under the *nlg-1* promoter, and the red signal corresponds to DiI staning in amphid sensory neurons. There is no neuroligin expression in the DiI stained ADL, ASH, ASI, ASJ, ASK, AWB amphid neurons. However, there is expression of YFP in ALA, URX and AVJ neurons. The box inset shows a detail of YFP and DiI staining in a subset of neurons. The neurons indicated with arrows and green labels correspond to positive neuroligin expression. The neurons indicated with head arrows and white labels correspond to amphid neurons that are DiI stained. **B**. Overlapping specific expression of neuroligin in a subset of dopaminergic neurons, ADEs, as well as the AIY interneuron. Representative images from the strain carrying the double promoter P*ttx-3::mCherry*/P*dat-1::mCherry* and P*nlg-1*::*yfp* arrays are shown. The top image corresponds to a merged composition between both green and red channels. Bottom panel shows detail of neurons where neuroligin is expressed in both ADE and AIY neurons (pointed by white arrows).

However, *nlg-1* is expressed in a subset of *eat-4* glutamatergic neuron classes **(Fig 5, Supplement 1)**. In this context it is noteworthy that glutamatergic signals are required for both foraging and feeding behaviours (Hills, Brockie et al. 2004, Dalliere, Bhatla et al. 2016). In addition, neuroligin expression was localized specifically in a subset of dopaminergic sensory neurons, ADEs (**Figs 5B)**, which also are involved in food-dependent behaviours (Hills, Brockie et al. 2004), as well as in the bilateral interneurons AIY (**Fig 5B**), that function to extend food-seeking periods (Shtonda and Avery 2006). Finally, by positional identification of cell bodies we found expression of *nlg-1* in the sensory neurons ALA and URX, as well as the interneuron AVJ (**Fig 5A**).

## DISCUSSION

The *C. elegans* neuroligin loss of function mutant is unable to maintain the same high rate of pharyngeal pumping in the presence of food compared to wild-type. This suggests that neuroligin is required in the circuitry that regulates feeding behaviour in response to sensory cues arising from bacteria. The activity of the pharynx is regulated by local and hormonal excitatory and inhibitory systems that converge to fine-tune food intake in a context-dependent manner (Dillon, Franks et al. 2015, Dalliere, Bhatla et al. 2016, Dillon, Holden-Dye et al. 2016). Insight into pharyngeal regulation came from a study in which it was shown that laser ablation of all pharyngeal neurons does not completely abolish pharyngeal pumping (Avery and Horvitz 1989). It has been proposed that there is an intrinsic myogenic rhythm that is modulated by the pharyngeal nervous system. This system is embedded underneath the pharyngeal basal membrane and is connected to the extrapharyngeal network via the RIP neurons (Bhatla, Droste et al. 2015). This provides a neural pathway for regulation of feeding behaviour by the extrapharyngeal nervous system (Trojanowski, Raizen et al. 2016) in response to sensory cues arising from food (Bhatla, Droste et al. 2015). However, ablation of RIP does not remove all food-dependent pharyngeal behaviours suggesting that other pathways, either intrinsic to the pharynx, for example the sensing of bacteria in the pharyngeal lumen, or arising through extrapharyngeal neurohormonal signals are important (Dalliere, Bhatla et al. 2016). The involvement of neuroligin in the first mechanism is unlikely as there is no evidence for *nlg-1* expression in the pharynx. This suggests neuroligin is required in an extrapharyngeal circuit for robust up-regulation of pharyngeal pumping in the presence of food.

Our data suggest that neuroligin-dependent extrapharyngeal processing of the food cue modifies the manner in which the pharynx responds to the presence of food (**Fig 6**) and that it is required to decrease the duration of a pharyngeal pump, most likely by increasing activity of the glutamatergic neuron M3. An inability to execute this control in the presence of food is consistent with the observation that *nlg-1* is unable to sustain a high level of pumping in the presence of food as a long pump duration is incompatible with fast pumping. We considered where in the extrapharyngeal circuitry neuroligin may mediate this effect by mapping the expression pattern of *nlg-1*. It may function in circuits detecting food-related stimuli from the environment through sensory neurons such us ALA and URX and those processing information through interneurons such us AIY and AUA (Coates and de Bono 2002, Chalasani, Chronis et al. 2007). In addition, our data show a co-expression of neuroligin in extrapharyngeal glutamatergic neurons and this mirrors neuroligin control of glutamate transmission in mammals (Graf, Zhang et al. 2004, Budreck, Kwon et al. 2013). Neuroligin is also expressed in the ADE dopaminergic neurons and this could indicate a deficit in the dopaminergic contribution to food-related behaviours in *nlg-1* (Hills, Brockie et al. 2004). In this context it is interesting to note that in mice neuroligin-3 modulates inhibition onto dopaminergic neurons (Rothwell, Fuccillo et al. 2014).

**Figure 6.**
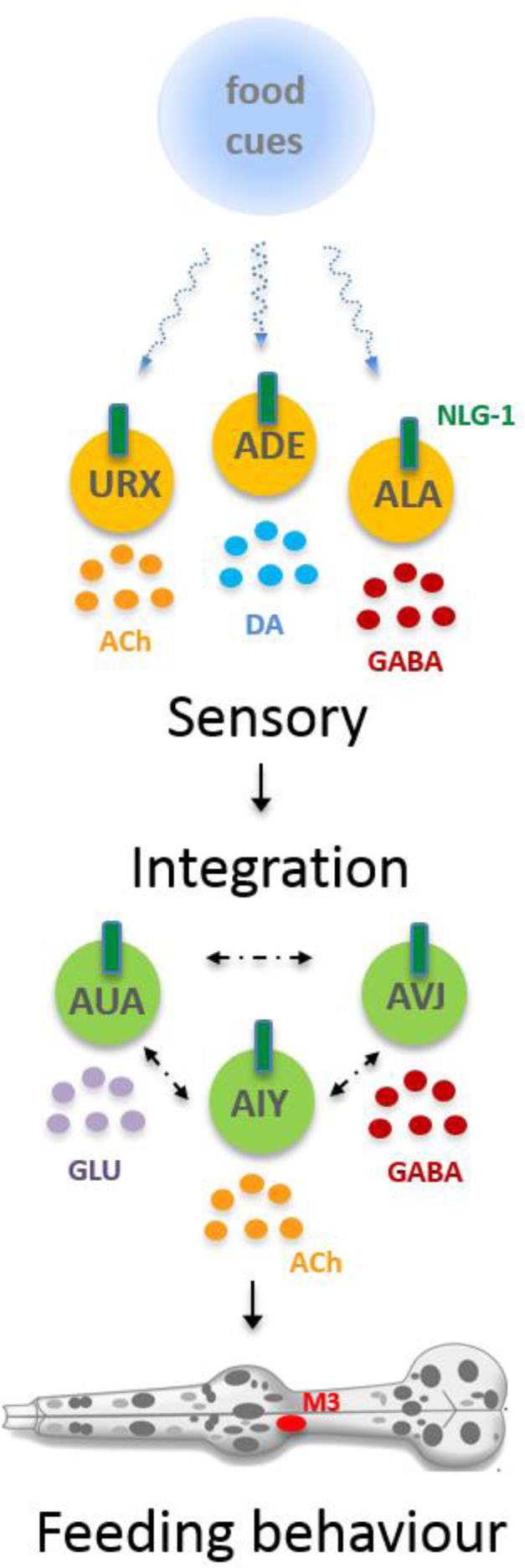
Hypothetical model where neuroligin expressing cells may act to sense and integrate food cues to up-regulate feeding. The proposed model show how neuroligin, located extrapharyngeally in a subset of neurons in the nervous system, is involved in the regulation of feeding-dependent behaviours, sensing cues from food, through communication between sensory neurons and interneurons by modulating pharynx activity in response of food cues by repolarization of muscles in the metacorpus and isthmus. Thus, neuroligin controls the pharyngeal pumping through processing and integration of multiple convergent pathways of different nature: cholinergic, dopaminergic, glutamatergic and GABAergic. The neurotransmitter representation (in yellow, blue, red and violet) is based on the transmitter that each specific neuron releases. Neuroligin is located in a subset of identified sensory neurons (shown in yellow) in the head ganglia. In addition neuroligin is also found in a subset of interneurons (shown in green), including AUA and AIY. Neuroligin is also present in the sensory neuron ADE which releases dopamine (DA). ADE signals in a sensory axis involving touch sensation and pheromone sensation processing. ADE indirectly connects with the interneuron AUA which harbours dopamine receptors. AUA interneuron is involved in regulating social feeding behaviours along with URX neuron. Neuroligin is present in URX and ALA sensory neurons implicated in oxygen/CO_2_ sensation and mechanosensation, respectively. Finally, neuroligin is also present in the interneuron AIY controlling food and odor-evoked behaviours and food-seeking behaviours. This indicates that neuroligin can process, possibly jointly with other regulators, multiple aspects from food dependent stimuli to integrate information in order to generate a suitable food-dependent behaviour.

We observed *nlg-1* expression in ALA and AVJ which are GABA-positive cells (Gendrel, Atlas et al. 2016). This goes hand-in-hand with the electrophysiological data showing that GABA signalling deficient mutants phenocopy the neuroligin pharyngeal EPG phenotype. This indirectly suggests a pivotal role for regulation of GABAergic signalling by neuroligin which is required to sustain a high level of feeding in the presence of food. A role for neuroligin in controlling GABAergic signalling in the context of a food driven behaviour has parallels with recent evidence showing neuroligin organizes *C*. *elegans* GABAergic postsynapses (Maro, Gao et al. 2015, Tong, Hu et al. 2015, Tu, Pinan-Lucarre et al. 2015) and the neuroligin dependence of GABA cellular signalling at the body wall neuromuscular junction (Tu, Pinan-Lucarre et al. 2015).

In conclusion, we provide evidence for the altered processing of food cues in a neuroligin deficient mutant, *nlg-*1, that results in an inability to maintain a sustained level of feeding in the presence of an abundant food source. This altered processing of sensory cues imparted by neuroligin deficiency is interesting in the broader context of the role of neuroligin in human autism spectrum disorder which features dysfunctional sensory processing (Beker, Foxe et al. 2018). Our study delivers new insight into the way in which neuroligin is required for context-dependent behavioural responses (Calahorro, Alejandre et al. 2009, Hunter, Mullen et al. 2010, Calahorro and Ruiz-Rubio 2012). We suggest a model in which neuroligin functions in the extra-pharyngeal nervous system and links sensory detection of food to feeding behaviour through communication between sensory neurons and interneurons impacting in downstream pharyngeal circuits. Thus, neuroligin is required to organize the circuit(s) that integrate food sensory cues to generate an appropriate behavioural response.

## AUTHOR CONTRIBUTIONS

F.C. conceived, performed and interpreted experiments and co-wrote the manuscript; F.K. performed and analysed feeding behavioural experiments; J.D. helped generating transgenic lines; L.HD. with V.OC. designed the study, conceived and interpreted experiments and co-wrote the manuscript.

## ACKNOWLEDGEMENTS

We thanks James Rand for kindly sharing strains and unpublished data; Antonio Miranda-Vizuete for sharing the *trx-1*::*DsRed* strain. Additional strains were provided by the CGC, which is funded by NIH Office of Research Infrastructure Programs (P40 OD010440). This study was supported by a grant from Wessex Medical Trust - U.K. (V05) to F.C.

---

**Figure 4 Supplement 1.**
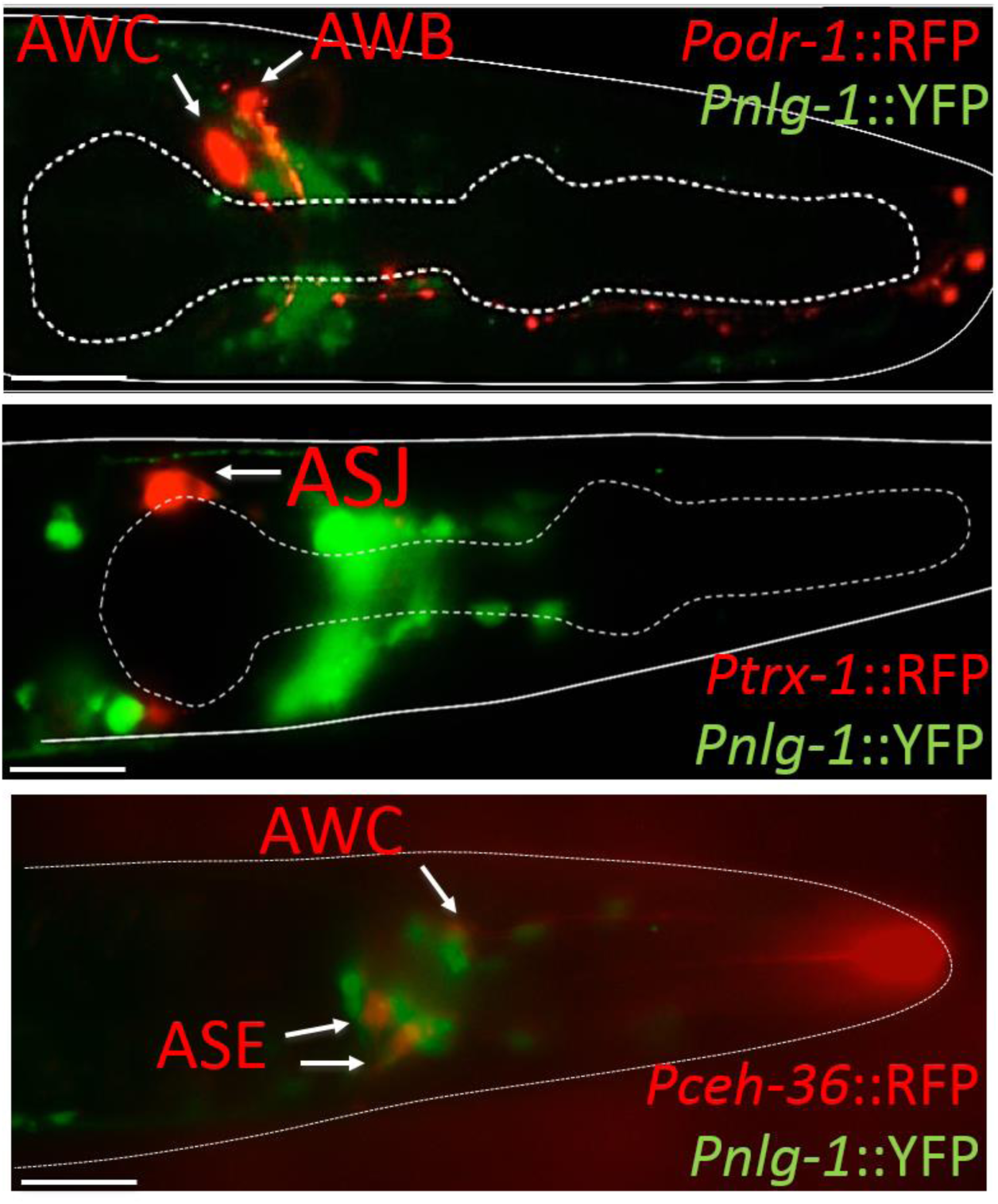
Co-localization experiments showing a non-expression of neuroligin in the main sensory neurons AWB, AWC, ASJ and ASE. A strain carrying a P*nlg-1*::YFP array was used to carry out genetic crosses with specifically labelled strains for different sensory neurons (see Material and Methods section for detail). A total of ten animals from each two independent crosses were examined. Representative merged images are shown. The scale bar indicates 50 μm.

**Figure 5 Supplement 1.**
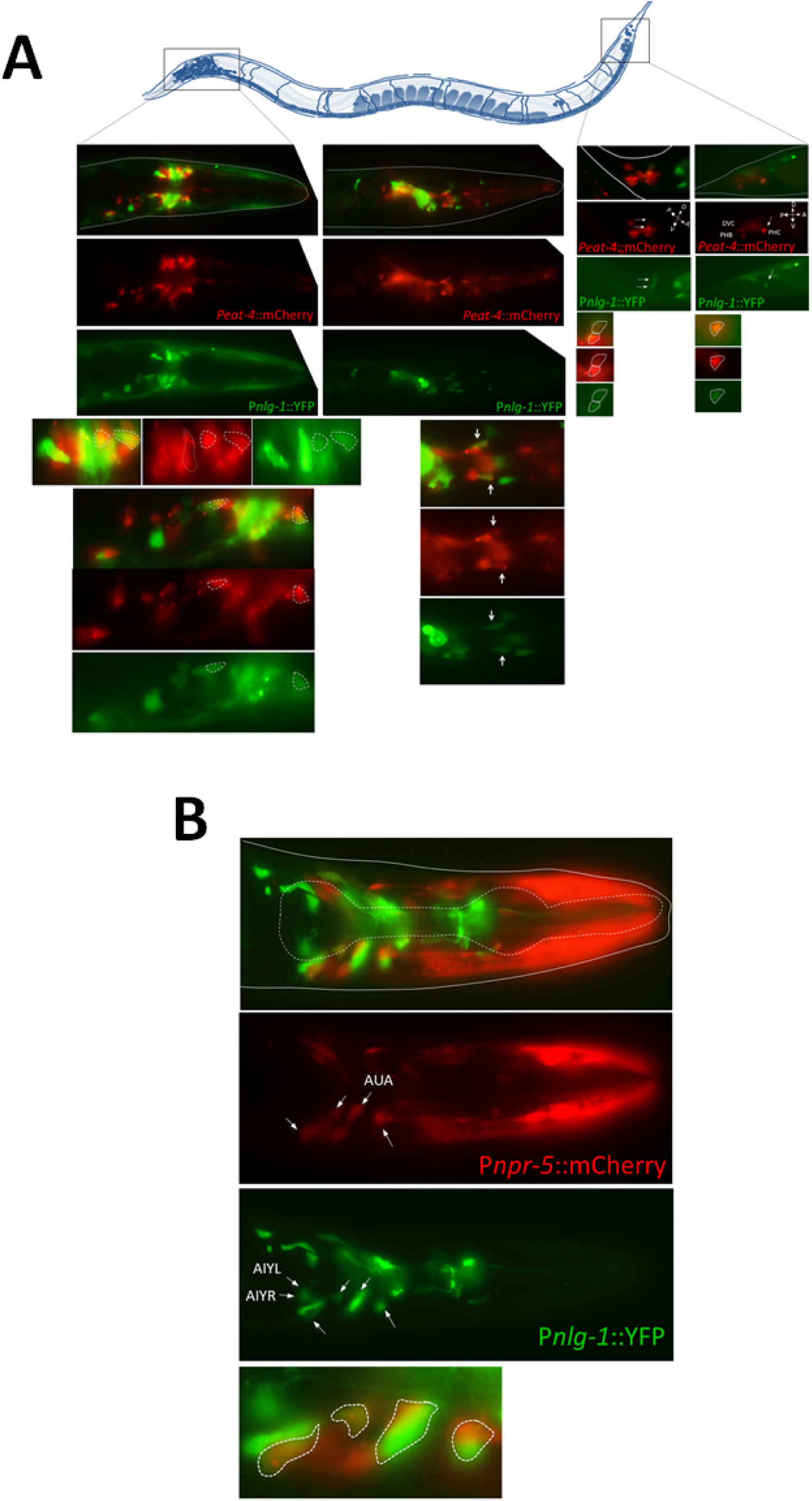
**A.** Overlapping expression of neuroligin in a subset of glutamatergic neurons expressing *eat-4*. Neuroligin is expressed in an unidentified subset of *eat-4* neurons in the head ganglia, as well as in the PHC phasmid neuron in the tail ganglia. The location of these ganglia is highlighted in the cartoon on the top of the panel by two boxes. Representative images from the strain carrying both arrays *Peat-4::mCherry*; P*nlg-1*::*yfp* in two diferent orientated planes are shown. The bottom panels correspond to detail view of *eat-4/nlg-1* matched expression. The white arrows point co-localized cell bodies. The dotted-line circles show co-localized cell bodies. The scale bar indicates 50 μm. **B.** The expression of neuroligin is matched to mCherry expression driven by the *npr-5* promoter. The *npr-5* promoter drives expression in a subset of sensory neurons (ADF, ASE, ASG, ASI, ASJ, ASK, AWA, AWB, IL2) and interneurons (AIA, AUA). Neuroligin is expressed in the AUA interneuron as well as in other unidentified subset of sensory neurons. A representative image from the strain carrying both arrays [P*npr-5::mCherry*]; P*nlg-1*::YFP are shown. The bottom panel shows detail of a subset of neurons where neuroligin is expressed. The white arrows point co-localized cell bodies. The dotted-line circles draw co-localized cell bodies.

**Supplementary Table 1.**
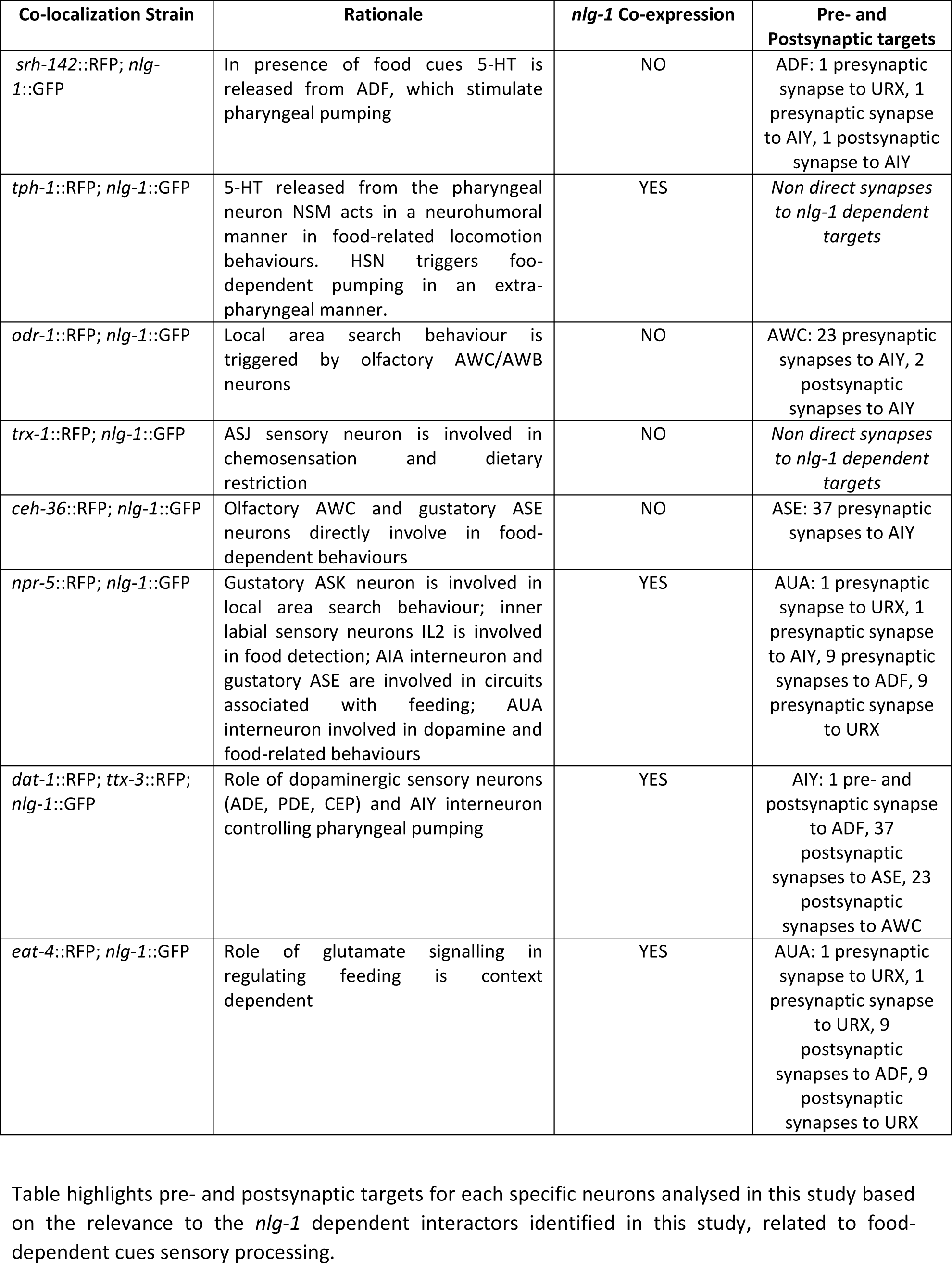
Results of co-localization experiments showing the strains used in this study, as well as the rationale to use specific markers labelling specific sensory and interneurons.

| Co-localization Strain | Rationale | *nlg-1* Co-expression | Pre- and Postsynaptic targets |
| --- | --- | --- | --- |
| *srh-142::RFP; nlg-1::GFP* | In presence of food cues 5-HT is released from ADF, which stimulate pharyngeal pumping | NO | ADF: 1 presynaptic synapse to URX, 1 presynaptic synapse to AIY, 1 postsynaptic synapse to AIY |
| *tph-1::RFP; nlg-1::GFP* | 5-HT released from the pharyngeal neuron NSM acts in a neurohumoral manner in food-related locomotion behaviours. HSN triggers food-dependent pumping in an extra-pharyngeal manner. | YES | *Non direct synapses to nlg-1 dependent targets* |
| *odr-1::RFP; nlg-1::GFP* | Local area search behaviour is triggered by olfactory AWC/AWB neurons | NO | AWC: 23 presynaptic synapses to AIY, 2 postsynaptic synapses to AIY |
| *trx-1::RFP; nlg-1::GFP* | ASJ sensory neuron is involved in chemosensation and dietary restriction | NO | *Non direct synapses to nlg-1 dependent targets* |
| *ceh-36::RFP; nlg-1::GFP* | Olfactory AWC and gustatory ASE neurons directly involve in food-dependent behaviours | NO | ASE: 37 presynaptic synapses to AIY |
| *npr-5::RFP; nlg-1::GFP* | Gustatory ASK neuron is involved in local area search behaviour; inner labial sensory neurons IL2 is involved in food detection; AIA interneuron and gustatory ASE are involved in circuits associated with feeding; AUA interneuron involved in dopamine and food-related behaviours | YES | AUA: 1 presynaptic synapse to URX, 1 presynaptic synapse to AIY, 9 presynaptic synapses to ADF, 9 presynaptic synapse to URX |
| *dat-1::RFP; ttx-3::RFP; nlg-1::GFP* | Role of dopaminergic sensory neurons (ADE, PDE, CEP) and AIY interneuron controlling pharyngeal pumping | YES | AIY: 1 pre- and postsynaptic synapse to ADF, 37 postsynaptic synapses to ASE, 23 postsynaptic synapses to AWC |
| *eat-4::RFP; nlg-1::GFP* | Role of glutamate signalling in regulating feeding is context dependent | YES | AUA: 1 presynaptic synapse to URX, 1 presynaptic synapse to URX, 9 postsynaptic synapses to ADF, 9 postsynaptic synapses to URX |

Table highlights pre- and postsynaptic targets for each specific neurons analysed in this study based on the relevance to the *nlg-1* dependent interactors identified in this study, related to food-dependent cues sensory processing.

## REFERENCES

Albertson, D. G. and J. N. Thomson (1976). “The pharynx of Caenorhabditis elegans.” Philos Trans R Soc Lond B Biol Sci 275(938): 299–325.

Avery, L. (1993). “The genetics of feeding in Caenorhabditis elegans.” Genetics 133(4): 897–917.

Avery, L. (1993a). “Motor neuron M3 controls pharyngeal muscle relaxation timing in Caenorhabditis elegans.” J Exp Biol 175: 283–297.

Avery, L. and H. R. Horvitz (1989). “Pharyngeal pumping continues after laser killing of the pharyngeal nervous system of C. elegans.” Neuron 3(4): 473–485.

Avery, L., D. Raizen and S. Lockery (1995). “Electrophysiological methods.” Methods in Cell Biology, Vol 48 48: 251–269.

Barkus, C., S. B. McHugh, R. Sprengel, P. H. Seeburg, J. N. Rawlins and D. M. Bannerman (2010). “Hippocampal NMDA receptors and anxiety: at the interface between cognition and emotion.” Eur J Pharmacol 626(1): 49–56.

Beker, S., J. J. Foxe and S. Molholm (2018). “Ripe for solution: Delayed development of multisensory processing in autism and its remediation.” Neurosci Biobehav Rev 84: 182–192.

Bemben, M. A., S. L. Shipman, R. A. Nicoll and K. W. Roche (2015). “The cellular and molecular landscape of neuroligins.” Trends Neurosci 38(8): 496–505.

Bhatla, N., R. Droste, S. R. Sando, A. Huang and H. R. Horvitz (2015). “Distinct Neural Circuits Control Rhythm Inhibition and Spitting by the Myogenic Pharynx of C. elegans.” Curr Biol 25(16): 2075–2089.

Brenner, S. (1974). “The genetics of Caenorhabditis elegans.” Genetics 77(1): 71–94.

Budreck, E. C., O. B. Kwon, J. H. Jung, S. Baudouin, A. Thommen, H. S. Kim, Y. Fukazawa, H. Harada, K. Tabuchi, R. Shigemoto, P. Scheiffele and J. H. Kim (2013). “Neuroligin-1 controls synaptic abundance of NMDA-type glutamate receptors through extracellular coupling.” Proc Natl Acad Sci U S A 110(2): 725–730.

Budreck, E. C. and P. Scheiffele (2007). “Neuroligin-3 is a neuronal adhesion protein at GABAergic and glutamatergic synapses.” Eur J Neurosci 26(7): 1738–1748.

Calahorro, F. (2014). “Conserved and divergent processing of neuroligin and neurexin genes: from the nematode C. elegans to human.” Invert Neurosci 14(2): 79–90.

Calahorro, F., E. Alejandre and M. Ruiz-Rubio (2009). “Osmotic avoidance in Caenorhabditis elegans: synaptic function of two genes, orthologues of human NRXN1 and NLGN1, as candidates for autism.” J Vis Exp(34).

Calahorro, F., L. Holden-Dye and V. O’Connor (2015). “Analysis of splice variants for the C. elegans orthologue of human neuroligin reveals a developmentally regulated transcript.” Gene Expr Patterns 17(2): 69–78.

Calahorro, F. and M. Ruiz-Rubio (2012). “Functional phenotypic rescue of Caenorhabditis elegans neuroligin-deficient mutants by the human and rat NLGN1 genes.” PLoS One 7(6): e39277.

Calahorro, F. and M. Ruiz-Rubio (2012). “Functional Phenotypic Rescue of Caenorhabditis elegans Neuroligin-Deficient Mutants by the Human and Rat NLGN1 Genes.” Plos One 7(6).

Chalasani, S. H., N. Chronis, M. Tsunozaki, J. M. Gray, D. Ramot, M. B. Goodman and C. I. Bargmann (2007). “Dissecting a circuit for olfactory behaviour in Caenorhabditis elegans.” Nature 450(7166): 63–+.

Chang, S., R. J. Johnston, Jr. and O. Hobert (2003). “A transcriptional regulatory cascade that controls left/right asymmetry in chemosensory neurons of C. elegans.” Genes Dev 17(17): 2123–2137.

Coates, J. C. and M. de Bono (2002). “Antagonistic pathways in neurons exposed to body fluid regulate social feeding in Caenorhabditis elegans.” Nature 419(6910):925–929.

Cohen, M., V. Reale, B. Olofsson, A. Knights, P. Evans and M. de Bono (2009). “Coordinated regulation of foraging and metabolism in C. elegans by RFamide neuropeptide signaling.” Cell Metab 9(4): 375–385.

Cook, A., C. J. Franks and L. Holden-Dye (2006). “Electrophysiological recordings from the pharynx.” WormBook: 1–7.

Dalliere, N., N. Bhatla, Z. Luedtke, D. K. Ma, J. Woolman, R. J. Walker, L. Holden-Dye and V. O’Connor (2016). “Multiple excitatory and inhibitory neural signals converge to fine-tune Caenorhabditis elegans feeding to food availability.” FASEB J 30(2): 836–848.

Dillon, J., I. Andrianakis, K. Bull, S. Glautier, V. O’Connor, L. Holden-Dye and C. James (2009). “AutoEPG: Software for the Analysis of Electrical Activity in the Microcircuit Underpinning Feeding Behaviour of Caenorhabditis elegans.” Plos One 4(12).

Dillon, J., C. J. Franks, C. Murray, R. J. Edwards, F. Calahorro, T. Ishihara, I. Katsura, L. Holden-Dye and V. O’Connor (2015). “Metabotropic Glutamate Receptors: MODULATORS OF CONTEXT-DEPENDENT FEEDING BEHAVIOUR IN C. ELEGANS.” J Biol Chem 290(24): 15052–15065.

Dillon, J., L. Holden-Dye, V. O’Connor and N. A. Hopper (2016). “Context-dependent regulation of feeding behaviour by the insulin receptor, DAF-2, in Caenorhabditis elegans.” Invert Neurosci 16(2): 4.

Donna G. Albertson, J. N. T. (1976). “The pharynx of <em>Caenorhabditis elegans</em>.” Philosophical Transactions of the Royal Society of London. B, Biological Sciences 275(938): 299–325.

Flames, N. and O. Hobert (2009). “Gene regulatory logic of dopamine neuron differentiation.” Nature 458(7240): 885–889.

Flavell, S. W., N. Pokala, E. Z. Macosko, D. R. Albrecht, J. Larsch and C. I. Bargmann (2013). “Serotonin and the neuropeptide PDF initiate and extend opposing behavioral states in C. elegans.” Cell 154(5): 1023–1035.

Franks, C. J., L. Holden-Dye, K. Bull, S. Luedtke and R. J. Walker (2006). “Anatomy, physiology and pharmacology of Caenorhabditis elegans pharynx: a model to define gene function in a simple neural system.” Invert Neurosci 6(3): 105–122.

Franks, C. J., C. Murray, D. Ogden, V. O’Connor and L. Holden-Dye (2009). “A comparison of electrically evoked and channel rhodopsin-evoked postsynaptic potentials in the pharyngeal system of Caenorhabditis elegans.” Invert Neurosci 9(1): 43–56.

Fu, Z. and S. Vicini (2009). “Neuroligin-2 accelerates GABAergic synapse maturation in cerebellar granule cells.” Mol Cell Neurosci 42(1): 45–55.

Gendrel, M., E. G. Atlas and O. Hobert (2016). “A cellular and regulatory map of the GABAergic nervous system of C. elegans.” Elife 5.

Graf, E. R., X. Zhang, S. X. Jin, M. W. Linhoff and A. M. Craig (2004). “Neurexins induce differentiation of GABA and glutamate postsynaptic specializations via neuroligins.” Cell 119(7): 1013–1026.

Greene, J. S., M. Brown, M. Dobosiewicz, I. G. Ishida, E. Z. Macosko, X. Zhang, R. A. Butcher, D. J. Cline, P. T. McGrath and C. I. Bargmann (2016). “Balancing selection shapes density-dependent foraging behaviour.” Nature 539(7628): 254–258.

Hills, T., P. J. Brockie and A. V. Maricq (2004). “Dopamine and glutamate control area-restricted search behavior in Caenorhabditis elegans.” J Neurosci 24(5): 1217–1225.

Horvitz, H. R., M. Chalfie, C. Trent, J. E. Sulston and P. D. Evans (1982). “Serotonin and octopamine in the nematode Caenorhabditis elegans.” Science 216(4549): 1012–1014.

Hu, C., J. Dillon, J. Kearn, C. Murray, V. O’Connor, L. Holden-Dye and H. Morgan (2013). “NeuroChip: a microfluidic electrophysiological device for genetic and chemical biology screening of Caenorhabditis elegans adult and larvae.” PLoS One 8(5): e64297.

Hu, Z., S. Hom, T. Kudze, X. J. Tong, S. Choi, G. Aramuni, W. Zhang and J. M. Kaplan (2012). “Neurexin and neuroligin mediate retrograde synaptic inhibition in C. elegans.” Science 337(6097): 980–984.

Hunter, J. W., G. P. Mullen, J. R. McManus, J. M. Heatherly, A. Duke and J. B. Rand (2010). “Neu roligin-deficient mutants of C. elegans have sensory processing deficits and are hypersensitive to oxidative stress and mercury toxicity.” Dis Model Mech 3(5-6): 366–376.

Iwanir, S., A. S. Brown, S. Nagy, D. Najjar, A. Kazakov, K. S. Lee, A. Zaslaver, E. Levine and D. Biron (2016). “Serotonin promotes exploitation in complex environments by accelerating decision-making.” BMC Biol 14: 9.

LeDoux, J. E. (2000). “Emotion circuits in the brain.” Annu Rev Neurosci 23: 155–184.

Lee, K. S., S. Iwanir, R. B. Kopito, M. Scholz, J. A. Calarco, D. Biron and E. Levine (2017). “Serotonindependent kinetics of feeding bursts underlie a graded response to food availability in C. elegans.” Nat Commun 8: 14221.

Li, Z. Y., Y. D. Li, Y. L. Yi, W. M. Huang, S. Yang, W. P. Niu, L. Zhang, Z. J. Xu, A. L. Qu, Z. X. Wu and T. Xu (2012). “Dissecting a central flip-flop circuit that integrates contradictory sensory cues in C. elegans feeding regulation.” Nature Communications 3.

Maro, G. S., S. Gao, A. M. Olechwier, W. L. Hung, M. Liu, E. Ozkan, M. Zhen and K. Shen (2015). “MADD-4/Punctin and Neurexin Organize C. elegans GABAergic Postsynapses through Neuroligin.” Neuron 86(6): 1420–1432.

Miranda-Vizuete, A., J. C. Fierro Gonzalez, G. Gahmon, J. Burghoorn, P. Navas and P. Swoboda (2006). “Lifespan decrease in a Caenorhabditis elegans mutant lacking TRX-1, a thioredoxin expressed in ASJ sensory neurons.” FEBS Lett 580(2): 484–490.

Molin, L., H. Schnabel, T. Kaletta, R. Feichtinger, I. A. Hope and R. Schnabel (1999). “Complexity of developmental control: analysis of embryonic cell lineage specification in Caenorhabditis elegans using pes-1 as an early marker.” Genetics 151(1): 131–141.

Niacaris, T. and L. Avery (2003). “Serotonin regulates repolarization of the C. elegans pharyngeal muscle.” J Exp Biol 206(Pt 2): 223–231.

O’Connor, R. M., B. C. Finger, P. J. Flor and J. F. Cryan (2010). “Metabotropic glutamate receptor 7: at the interface of cognition and emotion.” Eur J Pharmacol 639(1-3): 123–131.

Raizen, D. M. and L. Avery (1994). “Electrical-Activity and Behavior in the Pharynx of CaenorhabditisElegans.” Neuron 12(3): 483–495.

Raizen, D. M. and L. Avery (1994). “Electrical activity and behavior in the pharynx of Caenorhabditis elegans.” Neuron 12(3): 483–495.

Rogers, C. M., C. J. Franks, R. J. Walker, J. F. Burke and L. Holden-Dye (2001). “Regulation of the pharynx of Caenorhabditis elegans by 5-HT, octopamine, and FMRFamide-like neuropeptides.” J Neurobiol 49(3): 235–244.

Rothwell, P. E., M. V. Fuccillo, S. Maxeiner, S. J. Hayton, O. Gokce, B. K. Lim, S. C. Fowler, R. C. Malenka and T. C. Sudhof (2014). “Autism-Associated Neuroligin-3 Mutations Commonly Impair Striatal Circuits to Boost Repetitive Behaviors.” Cell 158(1): 198–212.

Sarafi-Reinach, T. R., T. Melkman, O. Hobert and P. Sengupta (2001). “The lin-11 LIM homeobox gene specifies olfactory and chemosensory neuron fates in C-elegans.” Development 128(17): 3269–3281.

Serrano-Saiz, E., R. J. Poole, T. Felton, F. Zhang, E. D. De La Cruz and O. Hobert (2013). “Modular control of glutamatergic neuronal identity in C. elegans by distinct homeodomain proteins.” Cell 155(3): 659–673.

Shtonda, B. B. and L. Avery (2006). “Dietary choice behavior in Caenorhabditis elegans.” J Exp Biol 209(Pt 1): 89–102.

Song, B. M., S. Faumont, S. Lockery and L. Avery (2013). “Recognition of familiar food activates feeding via an endocrine serotonin signal in Caenorhabditis elegans.” Elife 2: e00329.

Tanis, J. E., J. J. Moresco, R. A. Lindquist and M. R. Koelle (2008). “Regulation of serotonin biosynthesis by the G proteins Galphao and Galphaq controls serotonin signaling in Caenorhabditis elegans.” Genetics 178(1): 157–169.

Thutupalli, S., S. Uppaluri, G. W. Constable, S. A. Levin, H. A. Stone, C. E. Tarnita and C. P. Brangwynne (2017). “Farming and public goods production in Caenorhabditis elegans populations.” Proc Natl Acad Sci U S A 114(9): 2289–2294.

Tong, X. J., Z. Hu, Y. Liu, D. Anderson and J. M. Kaplan (2015). “A network of autism linked genes stabilizes two pools of synaptic GABA(A) receptors.” Elife 4: e09648.

Trojanowski, N. F., D. M. Raizen and C. Fang-Yen (2016). “Pharyngeal pumping in Caenorhabditis elegans depends on tonic and phasic signaling from the nervous system.” Sci Rep 6: 22940.

Tu, H., B. Pinan-Lucarre, T. Ji, M. Jospin and J. L. Bessereau (2015). “C. elegans Punctin Clusters GABA(A) Receptors via Neuroligin Binding and UNC-40/DCC Recruitment.” Neuron 86(6): 1407–1419.

Varoqueaux, F., G. Aramuni, R. L. Rawson, R. Mohrmann, M. Missler, K. Gottmann, W. Q. Zhang, T. C. Sudhof and N. Brose (2006). “Neuroligins determine synapse maturation and function.” Neuron 51(6): 741–754.

Xu, L., S. Choi, Y. Xie and J. Y. Sze (2015). “Cell-Autonomous Gbeta Signaling Defines Neuron-Specific Steady State Serotonin Synthesis in Caenorhabditis elegans.” PLoS Genet 11(9): e1005540.

